# A Systematic Review of the Ocean Acidification Research in India: Research Trends, Gaps and Recommendations

**DOI:** 10.64898/2026.01.30.702760

**Authors:** Amit Kumar, Kozhikkaparambil Kunjulakshmi, M.S. Silpa

## Abstract

Ocean acidification, a consequence of climate change, has become a significant threat to marine organisms. Globally, tremendous efforts have been made to understand its impact on different ecological and biological processes. In India, this research area is still not fully explored, but expanding at an exponential rate. Hence, it is essential to consolidate the fragmented knowledge into a systematic review, which will assist future researchers to develop their work. In this study, we utilized the Scopus, Web of Science and Ocean Acidification – International Coordination Centre bibliography to conduct a systematic review of ocean acidification research in India. We used the Biblioshiny package in R to conduct a bibliometric analysis, identify spatial and temporal research trends, and highlight the growth of literature in ocean acidification research, as well as existing knowledge gaps. We used the following keywords: ocean acidification, lowered pH, acidifying ocean, elevated carbon dioxide, elevated CO_2_, marine carbonate chemistry, shell decalcification and affiliation as India to obtain relevant publications. We selected 353 publications by applying relevance filtering and adherence to PRISMA guidelines. Almost one-third of the publications were non-primary articles. Among research articles, only 71 publications were found to investigate the response of marine organisms to ocean acidification. Majority of them involved single stressors, for a short term on very limited taxa. Lack of molecular-level investigation, multifactorial experimental design, and long-term observations were major gaps. This review aims to support researchers, policymakers, and other stakeholders involved in the planning, monitoring, and developing adaptation strategies. Finally, it provides recommendations for future research and policy development.

## 1. Introduction

Since the industrial revolution, human activities have injected approximately 750 petagrams of carbon into the atmosphere. The ocean acts as a major sink, absorbing around 26% of this atmospheric carbon dioxide (Friedlingstein et al. 2024). When carbon dioxide dissolves in seawater, it decreases seawater pH and alters seawater chemistry, leading to Ocean acidification. Ocean acidification has been reported to significantly impact marine organisms, thereby affecting marine food chains and food webs, and it is now recognised as a growing global concern (Doney et al. 2009; Hoesung, et al.2023).

Over the last few decades, researchers worldwide have made significant progress in understanding the response of marine organisms to ocean acidification stressors. Global efforts have been made to summarize these impacts through meta-analysis (Kroeker et al. 2013; Martins Medeiros and Souza 2023). Additionally, bibliometric analysis and visualizations of global ocean acidification research have also been conducted (Sahoo and Pandey 2022, 2020). Though global studies have provided valuable information on ocean acidification, regional studies are essential to understand location specific responses, vulnerabilities, and ecosystem dynamics. India, with its extensive coastline of almost 11,098 kilometres, diverse and unique marine ecosystems, and huge socio-economic dependence on coastal and marine resources, requires a systematic evaluation of its ocean acidification research trends and gaps. Despite this, a comprehensive assessment of research efforts on ocean acidification is lacking.

Hence, in the present study, we performed a systematic review of the ocean acidification research in India. This study compiles and synthesizes existing literature on ocean acidification research in India, describes the methods and monitoring approaches adopted by Indian researchers, identifies spatial and temporal research trends, growth of literature, and highlights existing knowledge gaps. This review aims to support researchers, policymakers, and other stakeholders involved in the planning, monitoring, and developing adaptation strategies. Finally, it provides recommendations for future research and policy development.

## 2. Materials and Methods

The literature encompassed the period of 1991 - 2025. We searched the publications in the Web of Science (WoS) (https://www.webofscience.com/wos/woscc/basic-search) and Scopus (https://www.scopus.com/search/form.uri?display=basic#basic) databases, as well as in the ocean acidification – International Coordination Centre (OA-ICC) bibliography (https://www.zotero.org/groups/2199752/oa-icc), and OA-ICC portal for ocean acidification biological response data (https://oa-icc.ipsl.fr/) to conduct a systematic review on ocean acidification research in India. The criteria for including articles were determined by the availability of publications indexed in the WoS, Scopus and OA-ICC databases, the foundation of our dataset. We believe that the dataset is representative of the current scientific trends.

Scopus is among the largest curated abstract and citation databases with extensive coverage of journals, books, and conference proceedings (Baas et al. 2020). Web of Science is the world’s oldest, most widely used database of research publications and citations (Birkle et al. 2020). We considered full-text journal articles, reviews, book chapters, proceedings articles, and notes, published in the English language. Data mining from Scopus and Web of Science under ‘All fields’ search category. The survey was carried out on 10^th^ October 2025, using following key words: “Ocean acidification”, “lowered pH”, “acidifying ocean”, “elevated CO_2_ marine”, “carbonate chemistry”, “shell decalcification” AND India. We filtered the datasets with Indian affiliation. OA ICC bibliography on Zotero and the OA-ICC portal for ocean acidification biological response data contained more than 10,000 articles and 1,600 articles, respectively. The results of the OA-ICC bibliography were further searched in the Scopus or Web of Science search outcome. The non-index publications were excluded from analysis. The final metadataset from Scopus and WoS was used to generate, visualize, and analyze bibliometric networks using Bibliometrix software in R Studio Version 4.3.2 (http://www.bibliometrix.org/) (Aria and Cuccurullo 2017). Bibliometrix excluded the duplicates and generated a combined dataset of 1824 publications.

We applied the Preferred Reporting Items for Systematic Reviews and Meta-Analysis (PRISMA) framework to systematically review articles and identify research gaps that require further investigation (Kunjulakshmi and Prakash 2025). The systematic review followed the scheme presented in Fig.1. Records were initially identified through data searching, followed by screening and data extraction. We applied strict inclusion and exclusion criteria to identify the relevant publications. Articles that did not meet the eligibility criteria were excluded. The eligibility criteria involved ineligible documentation type, ineligible study design, ineligible outcome, and unavailable reference, as well as assessing the relevance of the remaining articles by reviewing their abstracts and contents. The final step involved selecting studies for inclusion in the review based on those that passed the eligibility assessment.

**Fig. 1.**
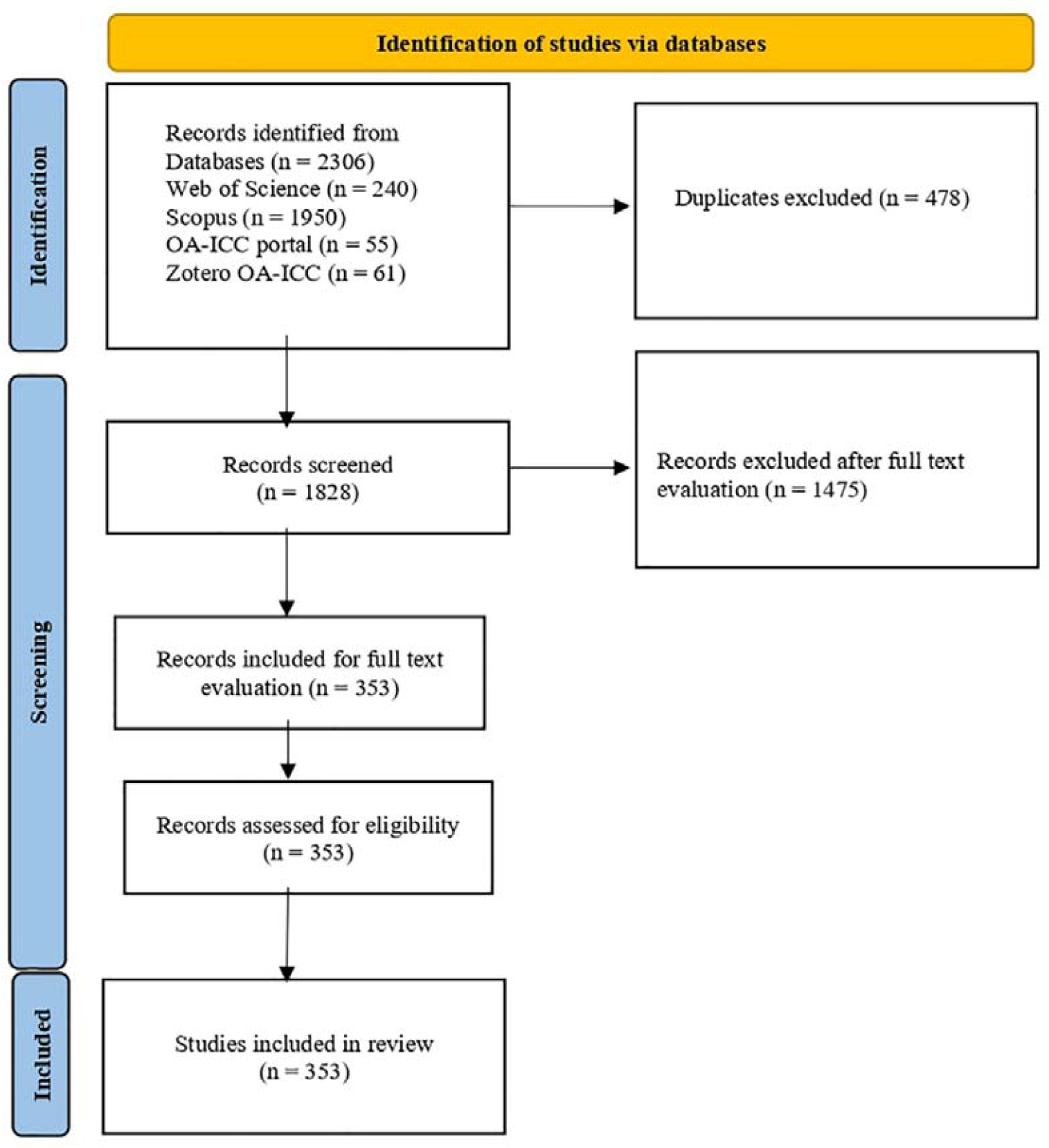
The Preferred Reporting Items for Systematic Reviews and Meta-Analyses (PRISMA) diagram used to search, screen, and select articles on Ocean acidification from India for the bibliometric review.

Since many of the keywords were common, such as CO_2_, pH, carbonate, decalcification, several irrelevant papers belonging to pure chemistry, nanoscience, mathematical modelling, biomedical science, etc, were extracted from databases. These papers were carefully examined through their title, abstract and text, and then excluded from the review *(N = 1471*). Finally, we were left with 353 publications, which were used for the construction of a network using Biblioshiny, a Shiny software that provides a web interface for Bibliometrix. The collected data was analysed and interpreted through bibliometric analysis, which allowed us to examine annual scientific production, scientific sources and their growth, the most cited articles, keyword analysis, trending topics as well as country-level collaboration networks.

## 3. Results

We obtained a total of 353 papers finally, after strict inclusion and exclusion criteria (Supplementary file S1). Our analysis suggests the research on ocean acidification picked up only in the last decade (Fig. 2). Though the time span in the search was 1991-2025, only 3 publications used ocean acidification as a keyword before 2010 (George et al. 1994; Sarma and Narvekar 2001; Peterson and Baringer 2009). These papers mostly deal with the carbon dynamics and biogeochemistry in the Arabian Sea and Bay of Bengal. Out of the total 353 publications, 107 (∼35%) were non-primary literature, comprising books, book chapters, editorial materials and reviews. A growth rate of almost 13% was found in ocean acidification publications.

**Fig. 2.**
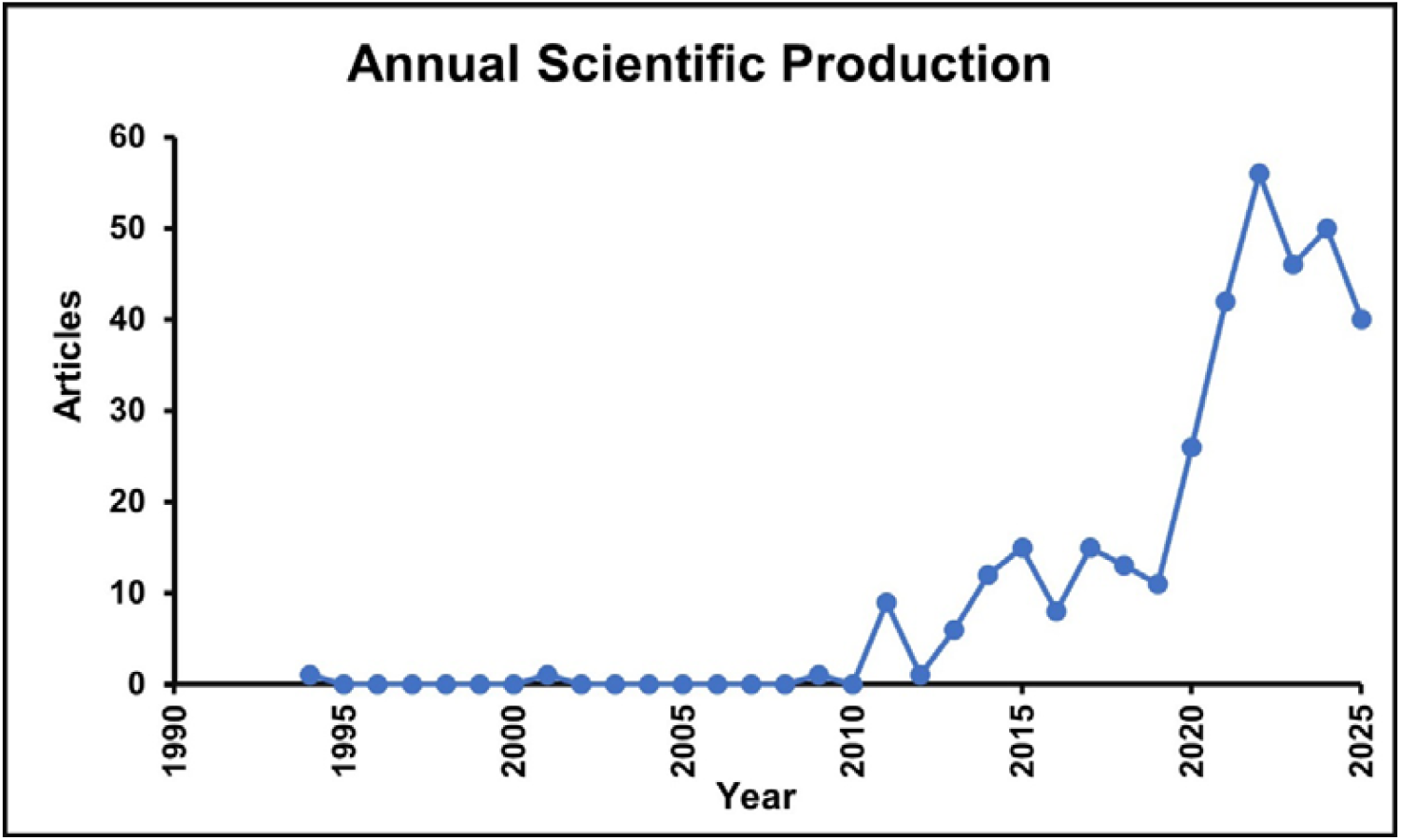
Number of research articles on ocean acidification published per year from 1991 – 10th October 2025.

We carefully evaluated the primary research articles and found that only 71 articles directly report the influence of ocean acidification or lowered pH conditions on the marine organisms. We found these studies were conducted on 18 taxa, with a maximum study on crustaceans (15 publications), followed by phytoplankton (14), macroalgae (9), and others (Table 1). More than 70% of the publications used lowered pH/ocean acidification as a single stressor, around 20% publications investigated ocean acidification and either ocean warming or nutrient or salinity as a second factor, and just 10% publications have considered a multifactorial experimental design combining ocean acidification, warming and a third factor, such as nutrient or oil pollution. Just over 15% of the studies have considered field-based observation, while the remaining 85% studies were conducted for a short term in laboratory conditions.

**Table 1:** Taxa representation and biological responses to ocean acidification. (Abbreviation: OA-Ocean acidification, OW-Ocean Warming).

| Taxa | Number of publications | OA | OA\|OW | Third factor | Remarks | Field-based observation | Lab-based |
| --- | --- | --- | --- | --- | --- | --- | --- |
| Bacteria | 5 | 3 | 1 | 1 | nutrient |  | 5 |
| Protists | 1 | 1 |  |  |  |  | 1 |
| Phytoplankton | 14 | 8 | 5 | 1 | 1 study used Dust induced OA | 3 onboard | 11 |
| Microalgae | 1 |  | 1 |  |  |  | 1 |
| Coccolithophores | 3 | 2 | 1 |  |  | 1 | 2 |
| Fleshy & Calcifying macroalgae | 9 | 8 |  | 1 | nutrient | 1 | 8 |
| Seagrasses | 1 | 1 |  |  |  | 1 |  |
| Mangroves | 1 | 1 |  |  |  | 1 |  |
| Macrobenthos | 1 | 1 |  |  |  | 1 |  |
| Zooplankton | 2 | 2 |  |  |  | 1 | 1 |
| Foramenifera | 1 | 1 |  |  | Salinity induced acidification |  | 1 |
| Coral | 2 |  | 2 |  |  |  | 2 |
| Cnidarians | 1 | 1 |  |  |  |  | 1 |
| Polychaete | 2 | 2 |  |  |  | 1 | 1 |
| Crustaceans | 15 | 9 | 4 | 2 | oil pollution | 1 | 14 |
| Mollusks | 6 | 5 |  | 1 | microplastic | 2 | 4 |
| Echinoderms | 2 | 2 |  |  |  | 1 | 1 |
| Fish | 4 | 2 | 1 | 1 | salinity |  | 4 |

Upon closer analysis of the response variables, for unicellular organisms, i.e. for bacteria, the studies have considered survivability, biosynthesis of extracellular polymeric substances, biofilm formation, and siderophore production (Patel, Shimpi, and Haldar 2020). For most of the studies on the autotrophs, including micro- and macro-algae, researchers have studied growth, photophysiology, primary and secondary metabolites and in a few cases oxidative status, in response to acidification (Vinuganesh, Kumar, Prakash, Alotaibi, et al. 2022; Vinuganesh, Kumar, Prakash, Korany, et al. 2022). For calcifiers, mainly the studies done on the crabs investigated the response of acidification on growth rate, survival, tissue biochemical constituents, and mineral content (Thangal et al. 2023, 2022). Similarly, among the fish, most studied response variables were physiological and oxidative stressors (Srinivasan et al. 2022).

In one study, Indian researchers have used pollutants to induce acidification to understand the response of phytoplankton (Sharma et al. 2022). Similarly, reduced salinity-induced acidification was used to investigate the response of foraminifera (Saraswat et al. 2015) (Table 1).

Our analysis revealed that ocean acidification research was published in 168 sources, including journals, books, and conference proceedings. Marine Pollution Bulletin published the highest number of 16 relevant publications, followed by Science of the Total Environment (14), and others (Supplementary Fig S1).

The majority of the research on ocean acidification is published by the Council of Scientific & Industrial Research (CSIR) institutions, such as the National Institute of Oceanography, Physical Research Laboratory, and Ministry of Earth Sciences (MoES) institutes, such as the Indian National Centre for Ocean Information Services, etc (Supplementary Fig. S2). In recent years, other institutions such as Sathyabama Institute of Science and Technology and others have begun working on the response of marine taxa to ocean acidification. Though more than 2300 researchers have authored/co-authored the ocean acidification papers, the author collaboration network revealed 6 groups working and publishing with a very low intergroup collaboration (Supplementary Fig. S3).

The analyses revealed ∼16% of the publications were done with global collaborations. Many of these collaborative publications are the outcome of research done by Indians during their PhD or postdoctoral research in foreign labs (Supplementary Fig. S4). Interestingly, only 3 molecular-level responses were published, and all of them were conducted in foreign labs on species not from Indian waters (Kumar et al. 2017; Stillman et al. 2020; Dineshram et al. 2021). Similarly, only 2 studies investigating coral responses to acidification were identified, and both were conducted in corals collected from the Caribbean Sea and work done in the USA (Guillermic et al. 2021; Eagle et al. 2022). Only one study on the Arctic diatom was performed in Germany by Indian researchers (Biswas 2022). Only two calcifying macroalgal responses have been evaluated, and both in South America (Korbee et al. 2014).

A thematic trend analysis examined the evolution of research topics over time (Fig. 3). It tracked the logarithmic frequency of different keywords and provided an overview of their temporal progression. The positioning of keywords on the right-side figure 3 indicates their increasing prominence in recent years. While earlier studies predominantly focused on keywords such as “warming”, “biofilm”, “carbon”, “macroalgae “and “acidification”, more recent research trends have shifted towards ocean acidification-related topics such as “ocean acidification”, “global warming”, “climate change”, “antioxidants”, “dissolved organic carbon”, “Indian ocean” underscoring an increasing study on ocean acidification in Indian ocean and multiple stressors experiments.

**Fig. 3.**
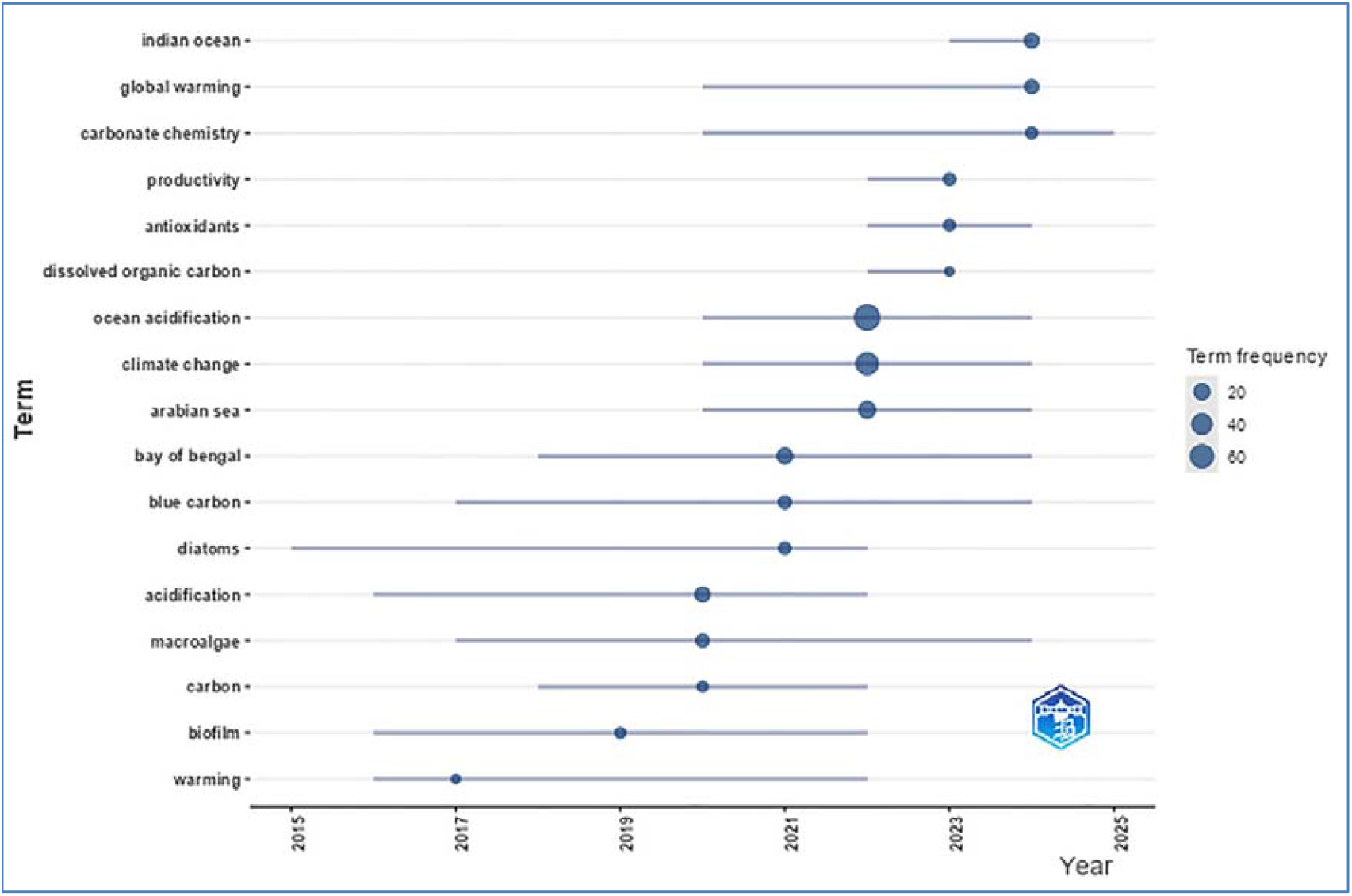
The trend topic of searched keywords in research articles.

The keyword co-occurrence analysis was used to identify the conceptual structure of ocean acidification studies and identify clusters that have shaped the field over time. By mapping and clustering terms extracted from keywords, the analysis provided insights into research trends and potential gaps in knowledge. Larger nodes in the co-occurrence network indicated higher keyword occurrences, while the links between nodes represented co-occurrences (i.e. keywords that frequently appear together). The thickness of these links signified the strength of the co-occurrence. In network visualization maps (Fig. 4), each label is denoted by a coloured node, with each colour representing a thematic cluster. The nodes and links within a cluster illustrate the tropical coverage and interconnections of research themes.

**Fig. 4.**
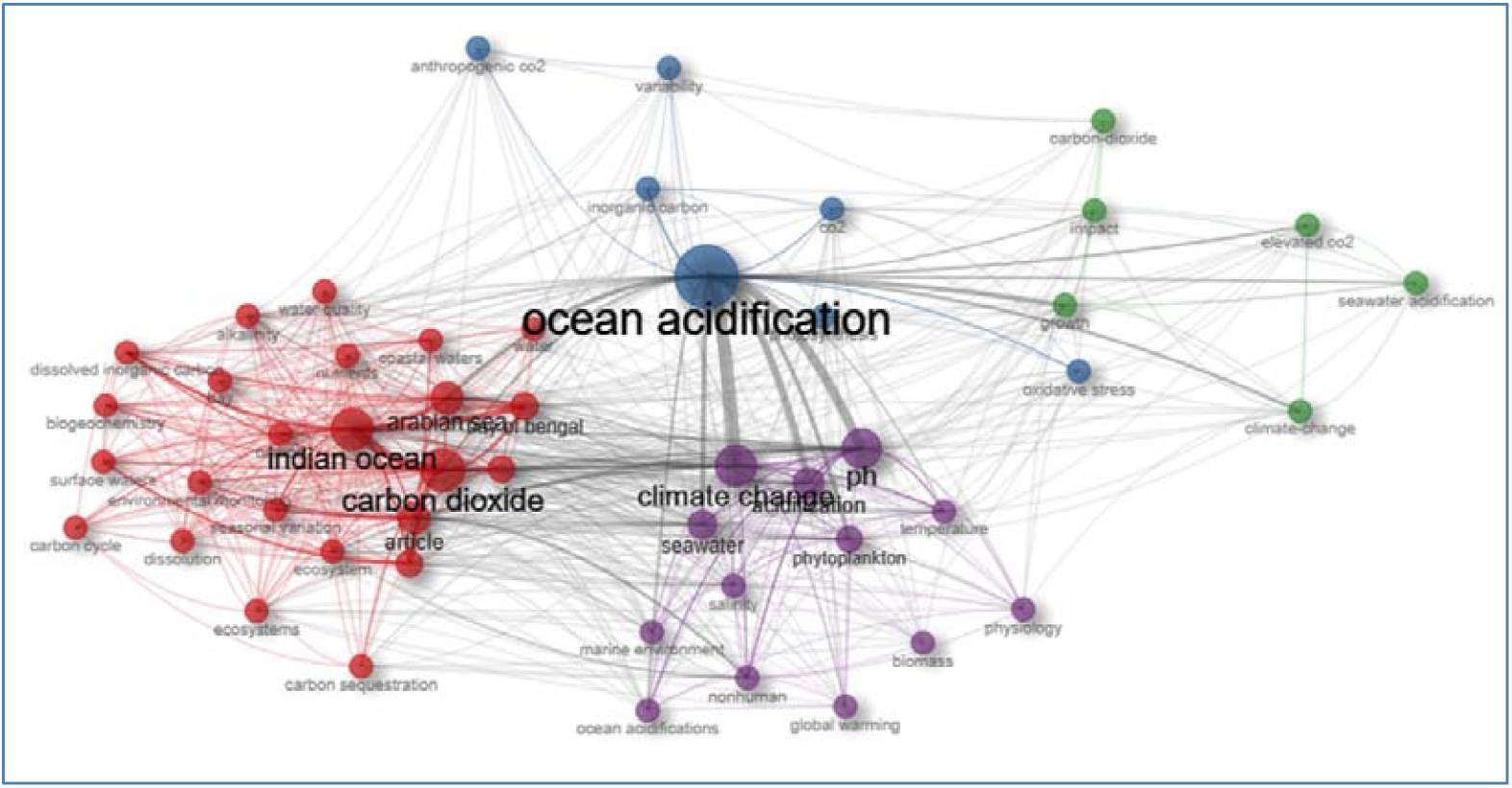
Keyword co-occurrence network map. A visualization of keyword relationships in ocean acidification research, where node size represents keyword frequency, edge thickness indicates co-occurrence strength, with thicker lines representing stronger associations between keywords. Different colours represent thematic clusters, grouping related research topics.

The clustering pattern reveals four major research foci: (1) mechanisms of ocean acidification and stress physiology, (2) broader climate change influences on marine systems, (3) regional carbon biogeochemistry of the Indian Ocean, and (4) organism-level impacts of elevated CO□. Cluster 1 includes keywords such as *‘*Ocean acidification,’ ‘CO□,’ ‘variability,’ ‘anthropogenic CO□,’ ‘inorganic carbon,’ ‘photosynthesis,’ and ‘oxidative stress’. This cluster highlights research examining how increased anthropogenic CO□ and changes in inorganic carbon chemistry influence photosynthetic activity and stress responses in marine organisms.

Cluster 2 encompasses *‘*Indian Ocean,’ ‘carbon dioxide,’ ‘Arabian Sea,’ ‘carbon sequestration,’ ‘seasonal variation,’ ‘carbon cycle,’ ‘alkalinity,’ ‘nutrients,’ ‘biogeochemistry,’ ‘dissolved inorganic carbon,’ ‘surface waters,’ ‘environmental monitoring,’ and ‘Bay of Bengal.’ This cluster represents a regionally focused theme centred on the carbon dynamics and biogeochemical processes of the Indian Ocean, with particular attention to spatial and seasonal variability, nutrient fluxes, and carbon storage potential.

Cluster 3 is composed of *‘*climate change,’ ‘pH,’ ‘global warming,’ ‘temperature,’ ‘marine environment,’ ‘salinity,’ ‘phytoplankton,’ ‘acidification,’ ‘physiology,’ and ‘biomass,’ suggesting an emphasis on climate-driven impacts on marine ecosystems. The inclusion of terms related to temperature, salinity, and phytoplankton indicates studies exploring how environmental fluctuations affect organismal physiology, productivity, and community-level responses.

Cluster 4 includes ‘carbon dioxide,’ ‘elevated CO□,’ ‘seawater acidification,’ ‘growth,’ ‘impact,’ and ‘climate change,’ reflecting studies concerned with experimental and impact-based assessments of elevated CO□ and acidification on marine organisms’ growth and development.

## 4. Discussion

### 4.1. A general research trend

We covered a time frame of the last 35 years; however, very few publications were found before 2010, indicating that attention to ocean acidification research has surged only in the past decade. There is a progressive evolution in the understanding of carbonate chemistry and CO□ dynamics in the North Indian Ocean over the past five decades. Early foundational work by Naqvi and Reddy (1979) marked the beginning of systematic chemical investigations in the open ocean of the Laccadive Sea in the Arabian Sea. By the early 1990s, attention had shifted toward the role of partial CO□ pressure in shaping carbonate system dynamics. Kumar et al. (1992) contributed significantly to this transition by examining factors influencing carbon components across the Arabian Sea. Initial research was primarily associated with the Joint Global Ocean Flux Study (JGOFS) project, focused on carbon dioxide fluxes, carbonate chemistry, and productivity patterns in the Arabian Sea (Krishnaswami and Nair 1996). The 58 dataset from this important project is available on the Pangaea database (https://www.pangaea.de/?q=author:email:ocean@csnio.ren.nic.in). The JGOFS was succeeded by the Bay of Bengal Process Studies (BOBPS), which was funded by the then Department of Ocean Development, Government of India, to investigate the CO_2_ air-sea exchange balance in the Bay of Bengal (PrasannaKumar et al., 2007). Sarma et al. (2002) incorporated a global perspective by using the World Ocean Circulation Experiment (WOCE) and the Geochemical Ocean Sections Study (GEOSEC) datasets to assess anthropogenic CO□ influences on aragonite dissolution in the Indian Ocean, marking a shift toward human-induced ocean acidification studies. Subsequent works, such as Bates et al. (2006), expanded temporal coverage by addressing seasonal fluctuations in Total CO□ and partial CO□ pressure in both the Arabian Sea and the Bay of Bengal.

Over the last few decades, there has been a tremendous increase in ocean observations through in situ buoys, Argo floats, scientific cruises, and long-term monitoring of coastal and marine waters along the east and west coasts of India (https://services.incois.gov.in/portal/datainfo/dip9700.jsp). India has been participating in international projects such as the International Ocean Carbon Coordination Project towards a sustainable global observation network for marine biogeochemistry (Bakker et al. 2014). More recently, (Chakraborty et al. 2024) summarized the changes in the Indian Ocean seawater pH in response to changes in the sea-surface temperature, salinity, and carbonate chemistry over the last 4 decades from 1989 to 2019, suggesting a decline of 0.014 - 0.015 units per decade. This study also highlighted the need to understand Indian Ocean acidification and identify its key drivers.

The increase in publications post 2010 coincides with a global rise in awareness of ocean acidification as a major stressor in the marine environment, following reports from the Intergovernmental Panel on Climate Change (IPCC) and other influential publications (Gattuso and Hansson 2011; Brewer 2013). Globally, research on ocean acidification has increased at a faster pace due to its predicted socio-economic impacts (Turley and Gattuso 2012). Though the number of publications from India has increased, it remains modest compared to developed nations such as the United States, Australia, the United Kingdom, Germany, China, other European countries or Canada (Sahoo and Pandey 2020). An important drawback of Indian publications is that one-third of them are non-primary literature, suggesting that experimental, hypothesis-driven studies are yet to mature. This also indicates that ocean acidification research is still in its infancy, with researchers consolidating the fragmented knowledge into review articles and book chapters. This may assist future researchers in developing their work.

### 4.2. Biological response and taxa representation

Eighteen taxa (Table 1), ranging from unicellular bacteria to autotrophs to fishes, were studied for their response to ocean acidification conditions. While studies on the diverse taxa look impressive, seven taxa, including protists, seagrass, mangroves, foraminifera, and sea anemones, were represented by one study each. This indicates biases in research focused on crustaceans and phytoplanktons, likely due to their ease of maintenance in laboratories, their availability for experiments, and their ecological and economic importance in the industrial sector. Lack of studies on many calcifying species, including native species of corals and calcifying macroalgae, is a significant concern. Expanding research to include benthic communities, habitat formers, and other calcifying species is essential for gaining a holistic ecosystem-level understanding on ocean acidification response. This pattern of narrow, largely centred on crustaceans, phytoplankton and macroalgae is opposite in comparison to global research, where the research encompasses a broad range of calcifiers, including corals, molluscs, and other marine organisms (Sahoo and Pandey 2020).

Moreover, most studies investigated a few selected response variables. Though these studies are very important, the lack of molecular-level studies limits our understanding of the mechanism behind species’ response to ocean acidification. Investigating molecular mechanisms involving transcriptomics, proteomics, and metabolomics is essential to uncover the adaptive response of species to acidification stressors. Additionally, a lack of studies involving organisms across multiple trophic levels, such as plant-herbivory interactions, restricts our ability to identify resilience mechanisms in the marine ecosystems of Indian waters. The limited number of studies focusing on field-based or in situ observations indicates that our understanding of ocean acidification response in the Indian context is inadequate.

The seas surrounding the Indian subcontinent – the Bay of Bengal and Arabian Sea are a complex basin due to unique bathymetry, hydrodynamics, and climatology. Factors such as monsoon systems and its consequences control the pH of seawater (Panchang and Ambokar 2021). Consequently, a few innovative studies used pollutants or salinity to reduce the seawater pH to observe species response. These context-specific studies are valuable but remain isolated examples. A recent experimental work by Shetye et al. (2020) demonstrated that the natural pH fluctuations can impact calcifying organisms such as sea urchins.

In recent years, long-term monitoring of seawater carbonate chemistry has been initiated at a few places, such as Sundarbans Biological Observatory Time Series (https://sites.google.com/view/sbots/home), which serves as a valuable resource (Land et al. 2019, 2015). However, such monitoring is sparse along the Indian coastline.

Due to the simplicity of the study design, most of the studies tested the response of organisms to single stressors. Globally, it has been recognized that organismal response to multiple stressor experiments may yield varying results. Responses can be additive, synergistic, or antagonistic (Rynkowski et al. 2025). However, the experimental setup, cost, and infrastructure could be major limitations, due to which not many studies have considered this approach in India.

### 4.3. Publication source, Research Connections and networking

The publication related to ocean acidification mainly appeared in the environmental and pollution-related journals (Supplementary Fig. S1), indicating that the field is often considered within the broad theme of anthropogenic stress and climate change. Most of the literature was published from CSIR and MoES institutes, which is understandable given their leadership in large collaborative projects in the past, such as the Joint Global Ocean Flux Study (JGOFS) and Bay of Bengal Process Studies (BOBPS), which were led by them (Supplementary Fig. S2). However, the field is gradually expanding beyond traditional oceanographic centres, with various academic institutions beginning to contribute.

The aggregation of a few author groups indicates a lack of inter-group collaborations (Supplementary Fig. S3). Such fragmentation restricts interdisciplinary knowledge transfer, which slows the progress of the field. To address weak international collaboration, it is essential to formulate formal bilateral and multinational research partnerships, and participate in global initiatives such as the Global Ocean Acidification Observing Network (GOA-ON) through the South Asia Regional Hub (https://www.goa-on.org/regional_hubs/saroa/about/introduction.php).

### 4.4. Limitations, gaps and way forward

Despite the Indian Ocean being the third-largest ocean basin in the world, there is a substantial knowledge gap about regional processes, trends, and ecosystem responses to acidification. It has historically been underrepresented in global open-access datasets. Indeed, the keyword distribution also suggested a strong emphasis on climate–carbon interactions, with relatively fewer studies addressing long-term adaptation, mitigation, and ecosystem-level responses, indicating potential research gaps in these areas. Lack of *in situ* and field-based studies in the pristine and critical marine habitats of India, such as the Lakshadweep archipelago, the Andaman and Nicobar Islands, the Gulf of Mannar, Palk Bay, Sundarbans, etc, is a major limitation. Most research conducted to date has been short-term, laboratory-based, which leaves a gap in understanding the response under complex, multifactorial stressors and natural settings. Additionally, no molecular-level studies on any Indian species are available.

Globally, attempts have been made to integrate socio-economic impact of ocean acidification; however, in the Indian context, besides mentioning ocean acidification in the blue economy policies, nothing much has been explored. Given the inclination of the Indian government towards the blue economy, estimating the impact of ocean acidification on marine resources is the need of the hour.

To advance ocean acidification research in India, we recommend establishing long term monitoring sites along the coast for measuring water carbon chemistry. We should aim to integrate with global networks such as GOA-ON. Mesocosm facilities should be built for long term-controlled experiments capable of simulating multi-stressors conditions to allow ecologically relevant studies. Restricted permits should be given to conduct research on critical taxa such as corals. In situ ecosystem-level studies on ecologically important coral reefs, seagrass meadows, seaweeds, mangroves and estuaries should be promoted. An omics-based approach should be promoted to underpin the molecular-level adaptive responses.

We emphasize the need to build an interdisciplinary team that includes marine biologists, economists, anthropologists, and social scientists, spanning research institutes, non-governmental organizations, and citizens to enhance the power and applicability of ocean acidification research.

## 5. Conclusion

Although ocean acidification research in India has begun, it is essential to accelerate the progress to achieve maturity in this field. To advance this field in India, it must enhance research infrastructure, expand funding opportunities, and promote capacity-building initiatives that empower early-career scientists to undertake interdisciplinary studies linking ocean chemistry, ecosystem responses, and climate change. India can greatly aid in comprehending and reducing the effects of climate change on marine environments by establishing a strong and internationally applicable framework for ocean acidification research with collective efforts.

## CRediT authorship contribution statement

**Amit Kumar:** Conceptualization, Data collection and Analysis, Investigation, Methodology, Visualization, Writing – first draft, review & editing, supervision; **Kunjulakshmi K:** Data collection and Analysis, Investigation, Methodology, Visualization, Writing-review and editing. **Silpa M S:** Data collection and Analysis, Visualization, Writing - review & editing.

## Acknowledgements

The authors would like to thank the management of Sathyabama Institute of Science and Technology for providing the research facilities to the Centre for Climate Change Studies.

## Data availability

All data used in the manuscript are presented in the form of graphs, tables, figures and supplementary files.

## Supplementary files

**Supplementary File S1**. List of publications used for the review on ocean acidification research from India

**Supplementary Fig. S1**. Most relevant source with publications on ocean acidification research from India.

**Supplementary Fig. S2**. Most relevant affiliations carrying out ocean acidification research in India

**Supplementary Fig.S3**. Authors collaboration on ocean acidification research in India

**Supplementary Fig. S4**. Global Research Collaboration Network. A world map depicting international research collaborations on ocean acidification. Countries are shaded in varying intensities of blue, with darker shades representing higher research output. Lines connecting countries indicate collaborative research networks, with thicker lines representing stronger collaborations between institutions.

