## Supplementary figures and images for "A Systematic Review of the Ocean Acidification Research in India: Research Trends, Gaps and Recommendations"

### Supplementary Fig S1

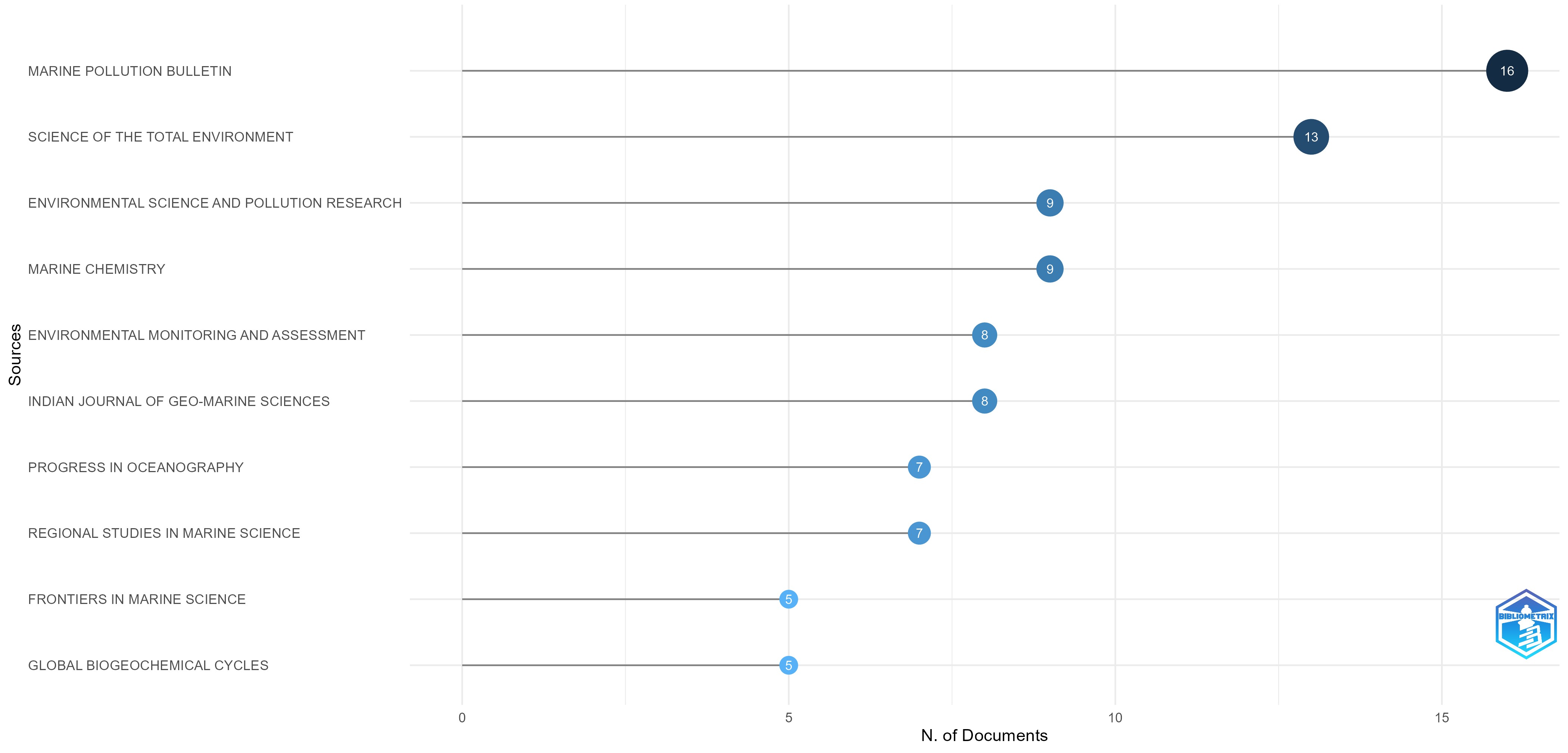

### Supplementary Fig S2

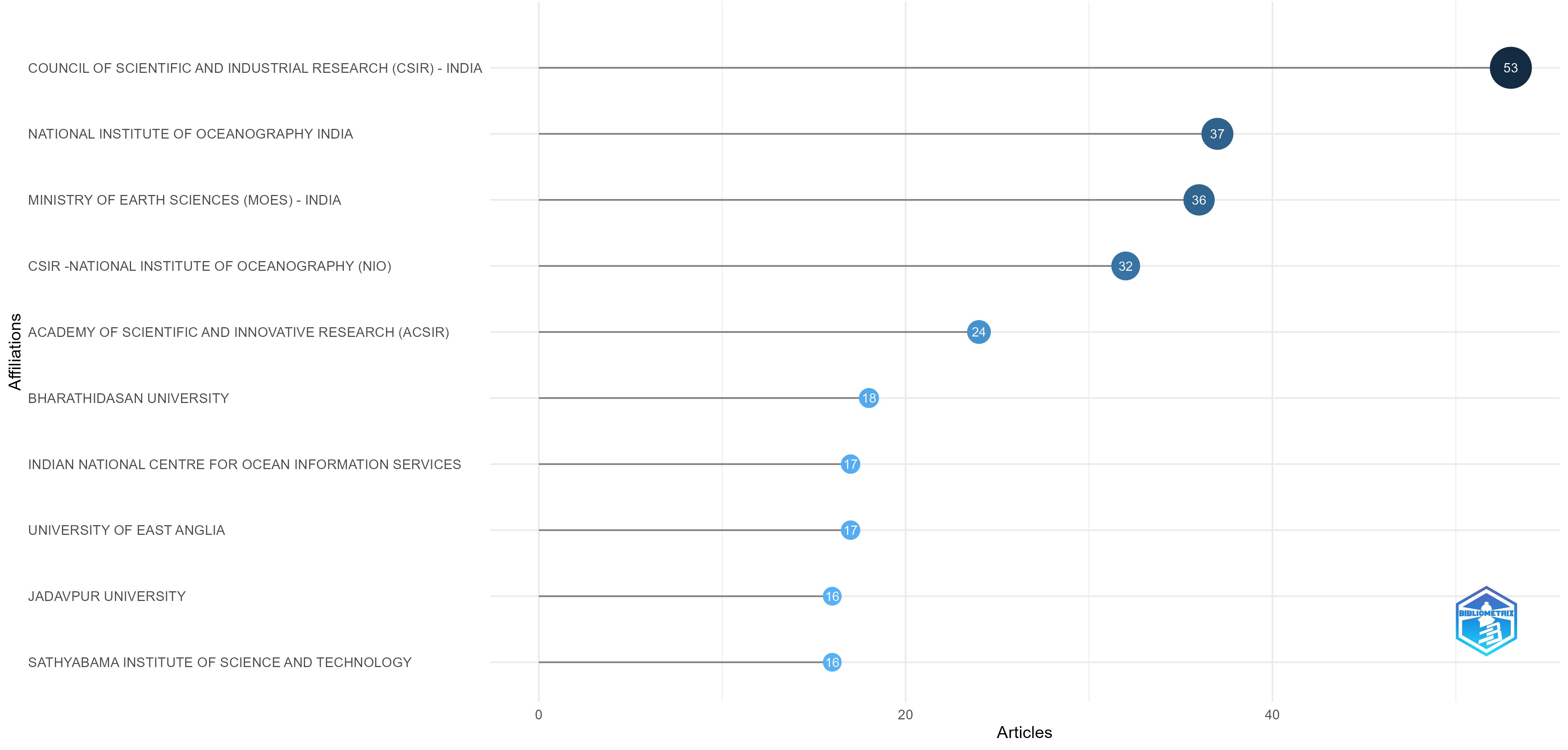

### Supplementary Fig S3

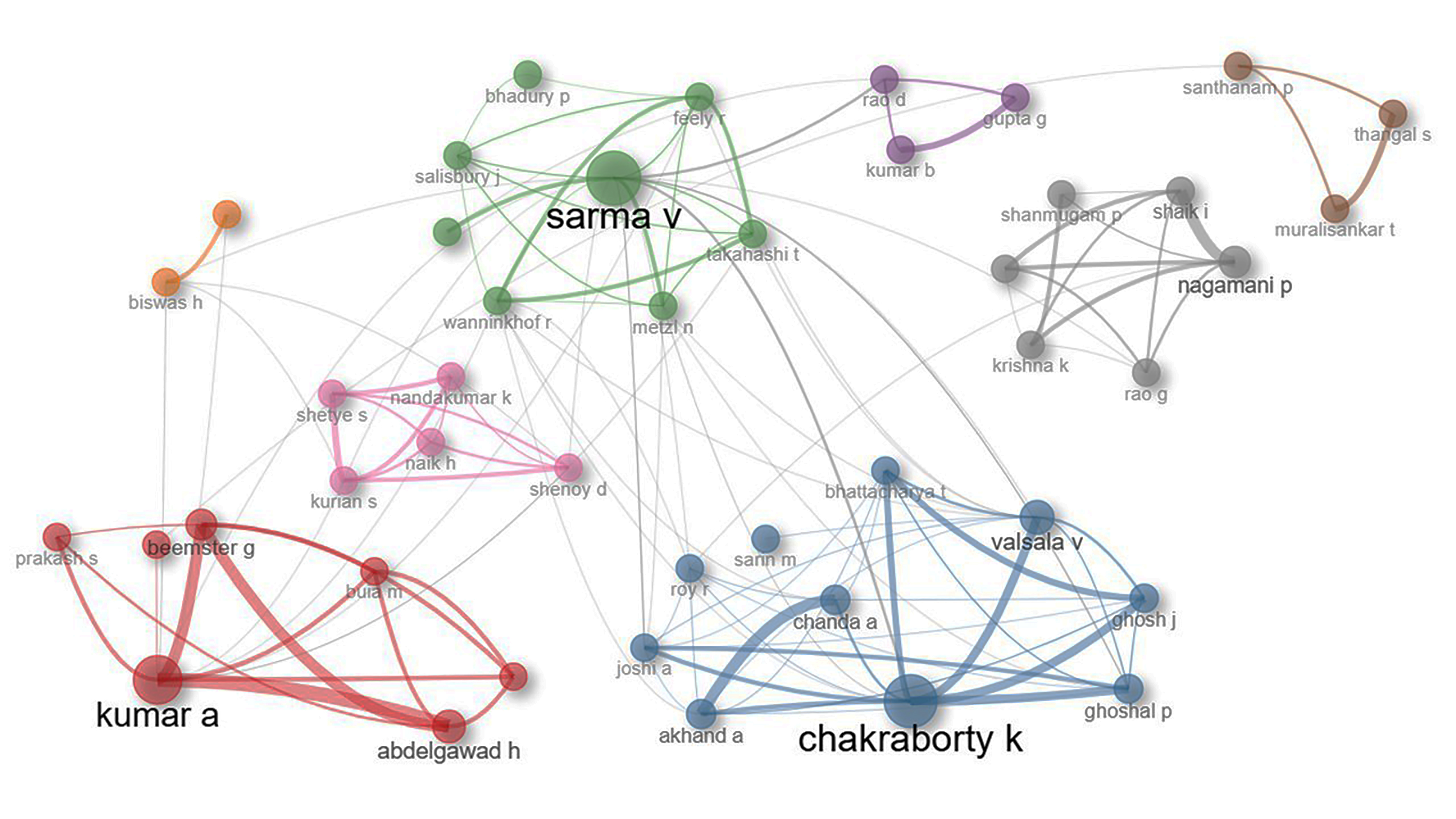

### Supplementary Fig S4

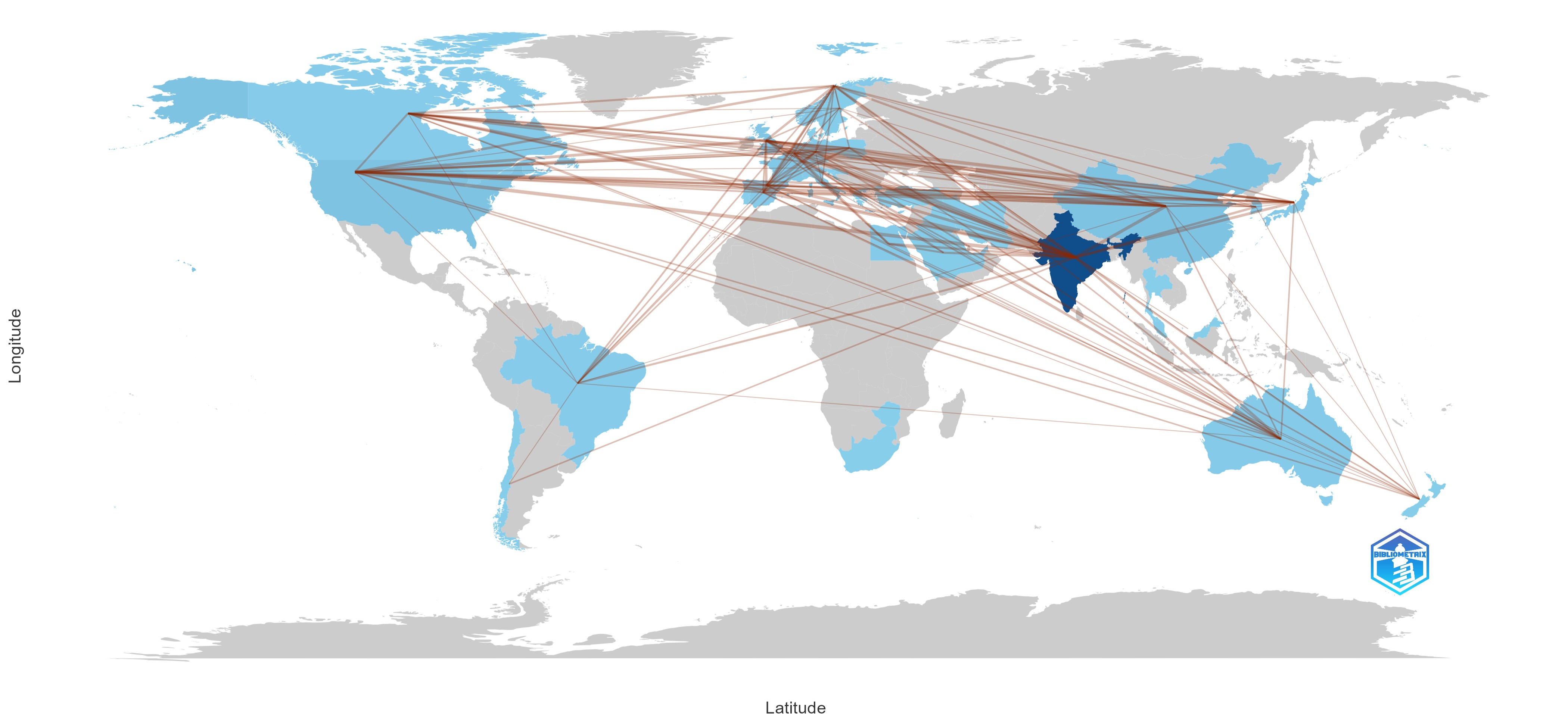
